# Rapid Phenotypic Detection of Carbapenem-Resistant *E. coli* with Fluorogenic Culture Media

**DOI:** 10.1101/2022.11.29.518369

**Authors:** Nodar Makharashvili, James O. McNamara

## Abstract

Faster, and simpler methods that determine the susceptibility of bacterial pathogens to antibiotics are needed to enable more effective treatment of bacterial infections and reduce the overuse of antibiotics. Here, we demonstrate a simple method for rapidly detecting bacteria and simultaneously determining their antibiotic susceptibility status. We demonstrate the method with *E. coli*, a high-impact pathogen, and meropenem, a member of the carbapenem class of antibiotics which is usually reserved for multi-drug resistant infections. Inclusion of a fluorogenic oligonucleotide substrate for endonuclease I, a well-conserved nuclease of the Enterobacteriaceae family, in a tryptic soy agar-based media enabled fluorescence-based detection of *E. coli* growth within 9 hours. Plating established carbapenem-resistant and -susceptible *E. coli* strains on this media with and without inclusion of meropenem, followed by fluorescent imaging yielded a clear phenotypic measure of the antibiotic susceptibility status of the strains in 9 hours. In addition to its simplicity and fast turnaround time, advantages of this approach include its ability to provide a measure of the bacterial load of the tested samples, and its easy integration into current microbiology laboratory workflows.

## Introduction

Optimal treatment of bacterial infections requires timely administration of antibiotics that are effective against the causative bacterial pathogen. Due to the potential for rapid progression of life-threatening infections such as bloodstream infections and endocarditis, the identification of effective antibiotics can be an urgent matter (Giacobbe et al., 2020). However, there is currently an unmet need for sufficiently rapid diagnostic methods that determine antibiotic susceptibilities of bacterial pathogens (Giacobbe et al., 2020). Current methods in common use require several steps and typically take 2 or more days from specimen collection to result, with turnaround times also depending on the type of infection (e.g., urinary tract infection, bloodstream infection, etc.).

Gram-negative species including various pathogens within the Enterobacteriaceae present an especially pressing need for such methods because they typically exhibit highly variable and unpredictable antibiotic susceptibility profiles (Livermore et al., 2011). Carbapenem-resistant Enterobacteriaceae (CRE) are a particularly problematic category of antibiotic resistant bacteria as carbapenems are usually reserved to treat multi-drug resistant infections (Papp-Wallace et al., 2011) and resistance to carbapenems is increasing within the Enterobacteriaceae (Livermore et al., 2011).

We previously developed a nuclease assay to rapidly detect the presence of Enterobacteriaceae (Flenker et al., 2017). This assay is based on a fluorogenic double-stranded oligonucleotide probe that is efficiently digested by endonuclease I, a well-conserved nuclease of the Enterobacteriaceae. We used this approach to rapidly detect bacteriuria in clinical urine specimens and have demonstrated its ability to detect various species of the Enterobacteriaceae family, including *E. coli, K. pneumoniae, P. mirabilis, S. marcescens*. Here, we incorporated an endonuclease I-responsive probe into agar media modified for optimal detection of this nuclease to enable detection of *E. coli* growth within 9 hours after inoculation. We additionally introduced meropenem into the media to enable detection of the growth of meropenem-resistant *E. coli* in 9 hours.

## Results

We adapted an approach we previously described for detecting various species of Enterobacteriaceae (Flenker et al., 2017) to enable rapid detection of the growth of these bacteria on solid culture media via their activation of an embedded fluorogenic oligonucleotide substrate. This approach provided the basis for a more rapid means of determining the antibiotic susceptibility of *E. coli* than alternatives such as chromogenic media. Initial evaluation carried out in a 96-well plate format (not shown) enabled clear differentiation of carbapenem resistant vs. susceptible *E. coli* strains that were applied to media with and without meropenem, with various bacterial concentrations tested.

Upon scaling up to 60 mm culture plates (**Figure 1)** fluorescence imaging revealed elevated fluorescence in several distinct spots on the media within 9 hours after application of dilute *E. coli* cultures. Note that the lower plate in the figure contains meropenem while the upper plate does not. Carbapenem-resistant, and susceptible *E. coli* strains were applied on the left and right sides of the plates, respectively. Fluorescent spots were visible on both sides of the upper (no meropenem) plate at 9 hours, but only on the left side of the meropenem-containing plate (i.e., the side where the carbapenem-resistant strain was applied). Taken together, this result indicates that the fluorogenic media optionally supplemented with an antibiotic can provide a rapid means of detecting *E. coli* (and likely related species) and also a phenotypic measure of the antibiotic susceptibility status of the bacteria.

**Figure 1.**
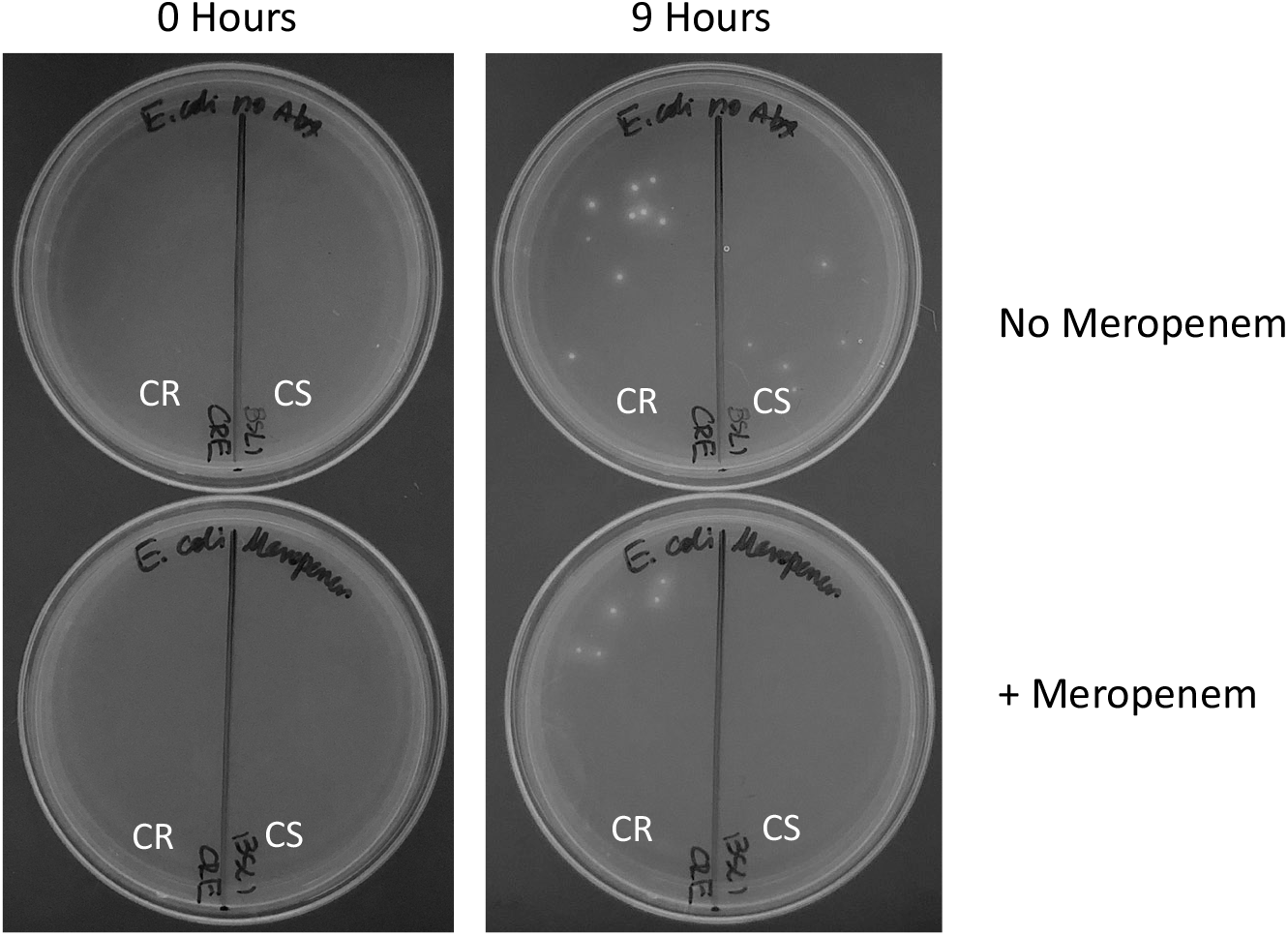
Selective detection of carbapenem-resistant *E. coli* with fluorogenic media containing an endonuclease I-responsive probe. Carbapenem -resistant (CR) and -sensitive (CS) *E. coli* strains were streaked on opposite sides of a plate with no meropenem (upper images) and a plate with meropenem (lower images) and the plates were incubated at 37 ⁰C. The fluorescent images shown were acquired at 0 and 9 hours after inoculation. Note the fluorescent spots visible on the 9-hour images for both strains on the non-selective plate, but only for the CR strain on the selective, meropenem-containing plate.

## Discussion

In the experiment presented here, we demonstrate phenotypic differentiation of low bacterial loads of meropenem-susceptible and -resistant *E. coli* strains with a novel fluorogenic culture media in 9 hours. This proof-of-concept demonstration could provide a technical foundation for addressing the unmet need for a rapid phenotypic test for carbapenem-resistant *E. coli* and other members of the Enterobacteriaceae. The simplicity of the test would seem to make it suitable for a variety of settings, including clinical microbiology laboratories and various low-resource settings.

Upon further development (see below), the approach described may provide a rapid CRE detection solution that could be integrated into clinical workflows in an almost identical manner to that of chromogenic media that is currently used for CRE screening of stool, rectal, and perianal swabs (Perry, 2017). Inexpensive and rapid identification of Enterobacteriaceae/CRE in positive blood cultures might also be enabled. In contrast to molecular tests in common use for determining antibiotic susceptibilities of Gram-negative isolates, the CRE fluorogenic media produces a phenotypic result. This is an important difference considering that molecular (genotypic) AST panels for Enterobacteriaceae are invariably incomplete (Giacobbe et al., 2020).

Further characterization of the method presented here with additional strains of both susceptible and resistant *E. coli* is needed to determine whether strain-dependent properties may impact results. Previous characterization of endonuclease I activity produced by various strains of *E. coli* suggest that its variability across different strains will be minimally impactful (Flenker et al., 2017). Moreover, the mechanism of meropenem resistance is not expected to affect performance because the method detects growth of resistant Enterobacteriaceae which will occur regardless of the resistance mechanism. Additional future objectives include defining the species inclusivity/exclusivity for the media, its limit-of-detection, and validation with clinical samples.

The concept for this test can likely be extended to yield rapid ASTs for additional bacterial species including other members of the Enterobacteriaceae such as *Klebsiella* spp., *Enterobacter* spp., *P. mirabilis, Serratia* spp., and Gram-positive pathogens such as *S. aureus* and *S. pseudintermedius*.

## Materials and Methods

A fluorogenic oligonucleotide probe that is a robust substrate for endonuclease I of *E. coli* was obtained via a custom order from Integrated DNA Technologies (IDT) of Coralville, IA. The probe was combined at 1 µM final concentration with tryptic soy agar (TSA) supplemented with NPT’s proprietary media additives, with or without 1.5 µg/ml meropenem as indicated in the figure. 2 ml of each fluorogenic media was poured per 60 mm culture plate. *E. coli* strains resistant (BAA-2452) or susceptible (ATCC 25922) to carbapenems were obtained from ATCC. Overnight cultures of the *E. coli* strains were grown in tryptic soy broth (TSB) in a shaking incubator at 37 ⁰C. Cultures were diluted 1:40 in fresh TSB, grown for an hour at 37 ⁰C and then further diluted in TSB to very dilute bacterial concentrations. 1 µL of each diluted culture was applied to one half of a plate with meropenem-containing fluorogenic media and to one half of a meropenem-free plate with a 1 µL calibrated loop. The plates were incubated at 37 ⁰C. The plates were imaged with a blue light transilluminator (IORodeo). Images were acquired at 0, 4, 6, 8, 9, and 24 hours after inoculation with an iPhone camera. Green channel images (shown in the figure) were isolated from the acquired color images with ImageJ.

